# *De novo* assembly of the whole genome of Moth bean (*Vigna aconitifolia*), an underutilized *Vigna* species of India

**DOI:** 10.1101/2023.05.18.540937

**Authors:** Sandhya Suranjika, Seema Pradhan, Rajwant K. Kalia, Nrisingha Dey

## Abstract

As a fast-growing legume species, moth bean (*Vigna aconitifolia)* has a unique habit of sustaining in conditions of higher temperatures and drought. This grain legume is also valued for it seeds which have one of the highest contents of proteins amongst all grain legumes. This plant can be a rich source of genomic resources, which can be applied to improve abiotic stress response in allied grain legumes and also help understand the biological processes governing their overall development. Here we generate a *de novo* genome assembly of *Vigna aconitifolia* using PacBio High-Fidelity reads and Hi-C sequencing data, with a total size of 409 Mb and contig N50 of more than 30Mb. We also annotated the genome for repeat sequences, found that the moth bean genome comprises of about 54% of repetitive sequences, and predicted 36950 protein-coding genes. Using the available RNA-Seq data for moth bean, we have developed a differential expression profile for various tissues of moth bean using the whole genome as a reference and identified simple sequence repeats that could be developed into viable molecular markers. This nascent study will provide insight into the identification of agronomically important genes and accelerate the genetic improvement of moth bean as well as other legume crops.

## Introduction

Moth bean (*Vigna aconitifolia* (Jacq.) Maréchal) is a non-conventional legume crop, belonging to the Fabaceae family and has a diploid (2n=2x=22) chromosome number. It is the most drought and heat-tolerant species of the Asian *Vigna* (Tomooka et al., 2011). Seeds of moth beans are appreciated due to high protein (22-24%) contents and carbohydrates associated with adequate amounts of minerals, vitamins, and some essential amino acids and unsaturated fatty acids (Tresina et al., 2017). Genetic diversity data and archaeological evidence suggest that Moth bean is native to India and cultivated mostly in some other countries including Bangladesh, Myanmar, China (Tomooka et al., 2011). Rajasthan (India’s driest state), is a significant moth bean producing state, accounting for over 86% of the country’s total area (Shah et al., 2019).

The genus *Vigna* consists of five subgenera and over 100 wild species. The subgenus *Ceratotropis*, also known as Asian *Vigna*, is a taxonomic group from which seven crops have been domesticated viz. moth bean (*Vigna aconitifolia* (Jacq.) Maréchal); minni payaru (*Vigna stipulacea Kuntze*); mung bean (*Vigna radiata* (L.) R. Wilczek); black gram (*Vigna mungo* (L.) Hepper); creole bean (*Vigna reflexo-pilosa* Hayata); rice bean (*Vigna umbellata* (Thunb.) Ohwi & Ohashi) and adzuki bean (*Vigna angularis* (Willd.) Ohwi & Ohashi) (Takahashi et al., 2016). Whole genome sequences have become available from the seven crops such as mung bean (Kang et al., 2014), black gram (Jegadeesan et al., 2021; Pootakham et al., 2021), rice bean (Guan et al., 2022), adzuki bean (Kang et al., 2015) whereas despite its economic potential, genomic resources for moth bean remain scarce.

Complete and precise reference genome assembly is indispensable for genetic and genome-wide studies of individual and multiple species. We have previously reported the *de novo* transcriptome assembly of *Vigna aconitifolia* var. RMO-435 (Suranjika et al., 2022), and we have now chosen the same variety for the whole genome assembly using PacBio long reads and high-throughput chromosome conformation capture (Hi-C) platform. The high-quality reference genome generated in this study aims to aid research on population genetic traits and functional gene discovery related to identify specific genes responsible for imparting important characteristics of moth bean.

## Materials and Methods

### Plant Materials and Growth Conditions

The seeds from a single plant of moth bean (*Vigna aconitifolia* var. RMO-435) were obtained from ICAR-CAZRI, Jodhpur, India (26° 15’ 49.9068’’ N, 73° 0’ 32.2452’’ E). They were grown in soil under aseptic conditions in the greenhouse (16 hr./8 hr. light/dark; 65% RH, 28° C ±2 temperature) after being germinated on a moist filter paper for 24 hrs. Young leaves were collected from the plants at five leaf stage, surface sterilized with 70% ethanol and stored at −80°C after freezing in liquid N_2_.

### Library preparation and sequencing

The preparation of libraries and their sequencing was outsourced to Nucleome Informatics Pvt Ltd, Hyderabad, India. Briefly, high quality genomic DNA (gDNA) was isolated from young leaves of moth bean and subjected to quality check; DNA purity ratios were checked with NanoDrop 2000 Spectrophotometer (Thermo Fisher Scientific, MA, USA) and Agarose gel electrophoresis (1% Agarose Gel) respectively. DNA was quantified using Qubit 3.0 Fluorometer (Thermo Fisher Scientific, MA, USA) and used to prepare HiFi SMRTBell libraries after shearing and size selection. The libraries were sequenced on the PacBio Sequel II platform to produce more than 400 Gb data.

Young leaves of moth bean were also used to construct Hi-C library after cross-linking with a buffer containing formaldehyde. The genomic DNA was then isolated from the leaves and digested with MboI restriction enzyme. The paired-end library was prepared (2X150) and sequenced on the NovaSeq platform (Illumina). The raw sequence data from both the platforms has been submitted to NCBI SRA database under the BioProject ID PRJNA967221.

### De novo genome assembly and quality assessment

The HiFi reads produced on the PacBio Sequel II platform were used to generate the primary assembly using Hifiasm software (Cheng et al., 2021). The primary unitigs (*.p.gfa) were used for further analysis. About 120Gb of sequence data generated from Hi-C sequencing was used for scaffolding the primary assembly using SALSA2 software (Ghurye et al., 2019). This gave the final assembly that was used for all the subsequent analyses in this study.

The final assembly was assessed for its quality by i) mapping the reads from RNA-seq library of the transcriptome of moth bean (Suranjika et al., 2022; PRJNA788336) and ii) using the BUSCO software package (Simão et al., 2015) to assess the percentage of single copy orthologs in the eukaryote (eukaryota_odb10) and fabales (fabales_odb10) databases.

### Genome annotation

We used the RepeatModeler software (http://www.repeatmasker.org/RepeatModeler/) to predict the repeat sequences in moth bean genome assembly. The predicted repeats were added to the repeat sequences identified in Fabids downloaded from the PlantRep database (http://www.plantrep.cn/) to form a single file. We then used the RepeatMasker software with default parameters (https://github.com/rmhubley/RepeatMasker/) to mask repeats in the moth bean genome. Genome was annotated using the RNA-Seq sequence reads of moth bean generated in a previous study (Suranjika et al., 2022). Briefly, the filtered, high quality reads from various tissues of moth bean were mapped onto the masked moth bean genome using HISAT2 (Kim et al., 2015). The output was converted to bam file and sorted before using it as an input parameter in the BRAKER2 pipeline (Li et al., 2009).

## Results

### De novo assembly of the moth bean genome

We assembled the genome of moth bean using the reads generated on PacBio Sequel II and NovaSeq 6000 platforms. A k-mer based analysis of the short reads using GenomeScope (https://github.com/schatzlab/genomescope) revealed that moth bean has a diploid genome with an estimated genome size of 360-380 Mb (Figure 1a). The high fidelity HiFi reads from PacBio platform were first assembled into a primary assembly containing 447 contigs, comprised of a total of 409 Mb of genome sequence. This assembly had an N50 value of 14.3 Mb with 54 contigs being more than 50Kb in length (Table 1). We then scaffolded the genome by integrating the Hi-C sequence data to get a final scaffold level assembly containing 340 scaffolds with an N50 value of 30.6 Mb. This assembly also displayed an improvement in other statistics. For example, the number of contigs with length more than 50Kb increased to 65 as compared to 54 in primary assembly (Table 1). Therefore, we used these scaffolds for repeat masking and annotation. We also observed that about 399Mb of the genome sequence was contained in 13 scaffolds while the rest 327 scaffold accounted for about only 10Mb of the genome (Supplementary data).

**Table 1.** Statistics for the whole genome assembly of Moth bean

| Assembly | Primary Assembly | After scaffolding with Hi-C |
| --- | --- | --- |
| Number of contigs/scaffolds | 447 | 340 |
| Largest contig/scaffolds | 58823228 bp | 58823228 |
| Total length | 409263063 | 409317563 |
| Number of contigs/scaffolds more than 50Kb in length | 54 | 65 |
| Length of contigs/scaffolds more than 50 Kb in length | 399447330 | 402100112 |
| **GC (%)** | 32.85 | 32.85 |
| **N50** | 14363921 | 30655968 |
| **L50** | 8 | 5 |
| **N's per 100 kbp** | 0 | 13.31 |

**Figure 1.**
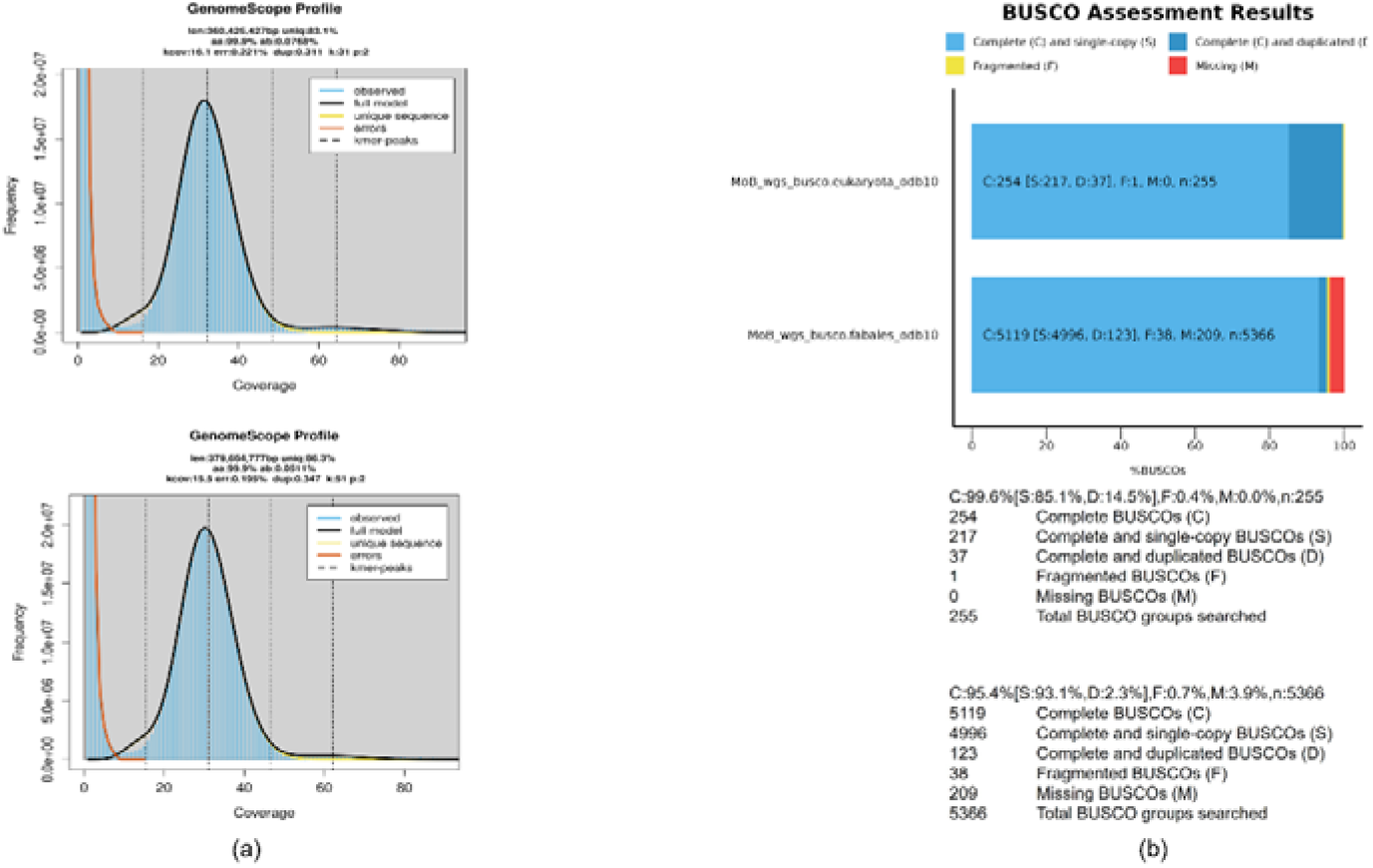
(a) k-mer based estimation of genome size for moth bean. (b) BUSCO assessment of genome assembly with eukaryota and fabales databases (MoB= **Mo**th **B**ean)

We mapped the filtered, paired end reads from the RNA-Seq data generated from various tissues of moth bean (Suranjika et al., 2022) onto the assemble genome and found that approximately 80% of the total reads could be mapped onto the genome. The genome assembly was also assessed with BUSCO where we found that 99% of complete BUSCOs in the eukaryote database and 95% of complete BUSCOs in the fabales database were represented in the moth bean genome (Figure 1b). All the parameters point towards a good quality genome assembly.

### Repeat masking and annotation

The repeat sequences were predicted in the genome as described earlier and we masked about 54% of the genome as repeat region. These included retroelements such as LINES, SINES, LTRs, DNA transposons and simple repeats (Table 2). We used the masked genome assembly to predict genes with the alignment information from RNA-Seq data of moth bean. The BRAKER2 software predicted 30075 genes consisting of 36950 coding sequences. These sequences were annotated using UniProt (https://www.uniprot.org/uniprotkb?query=reviewed:true), KEGG (https://www.genome.jp/tools/kaas/), KOG (https://mycocosm.jgi.doe.gov/help/kogbrowser.jsf) and Pfam (http://ftp.ebi.ac.uk/pub/databases/Pfam/current_release/) databases. A total of 36826 coding sequences found a match in these databases and could be assigned a functional role which gives a high percentage of annotation at 99% (Figure 2).

**Table 2.**
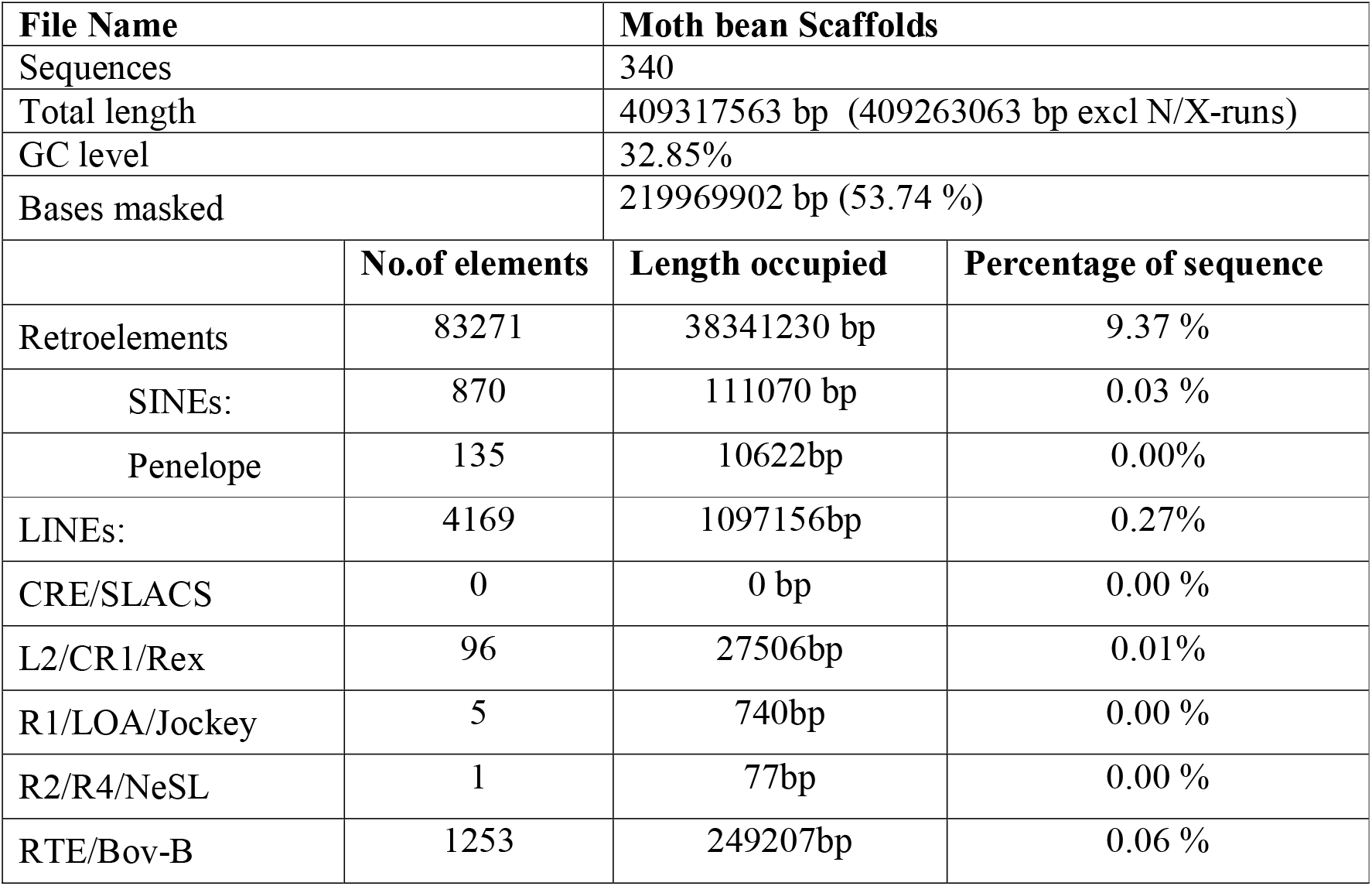
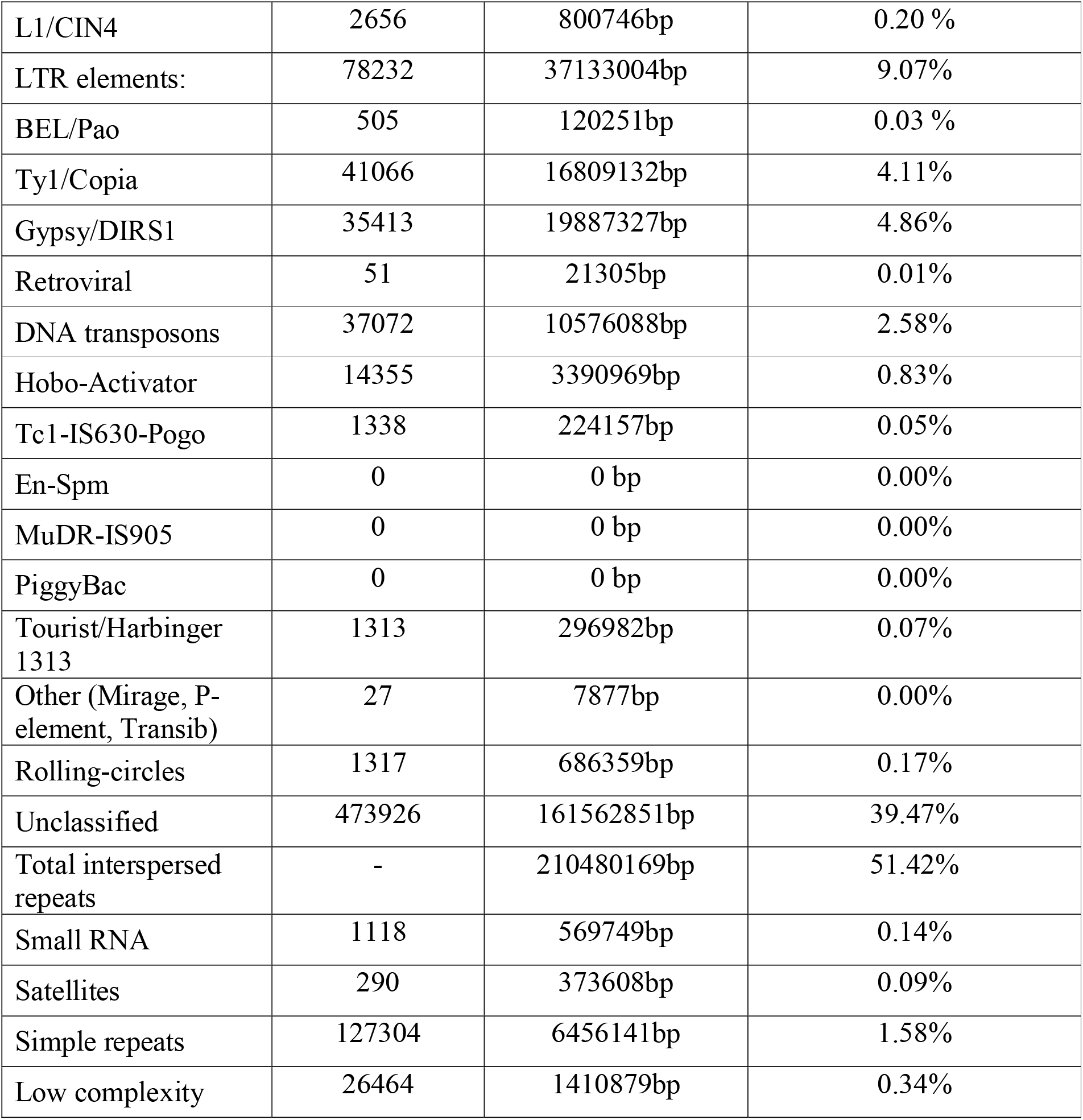
Repeat annotation in the genome of Moth bean

**Figure 2.**
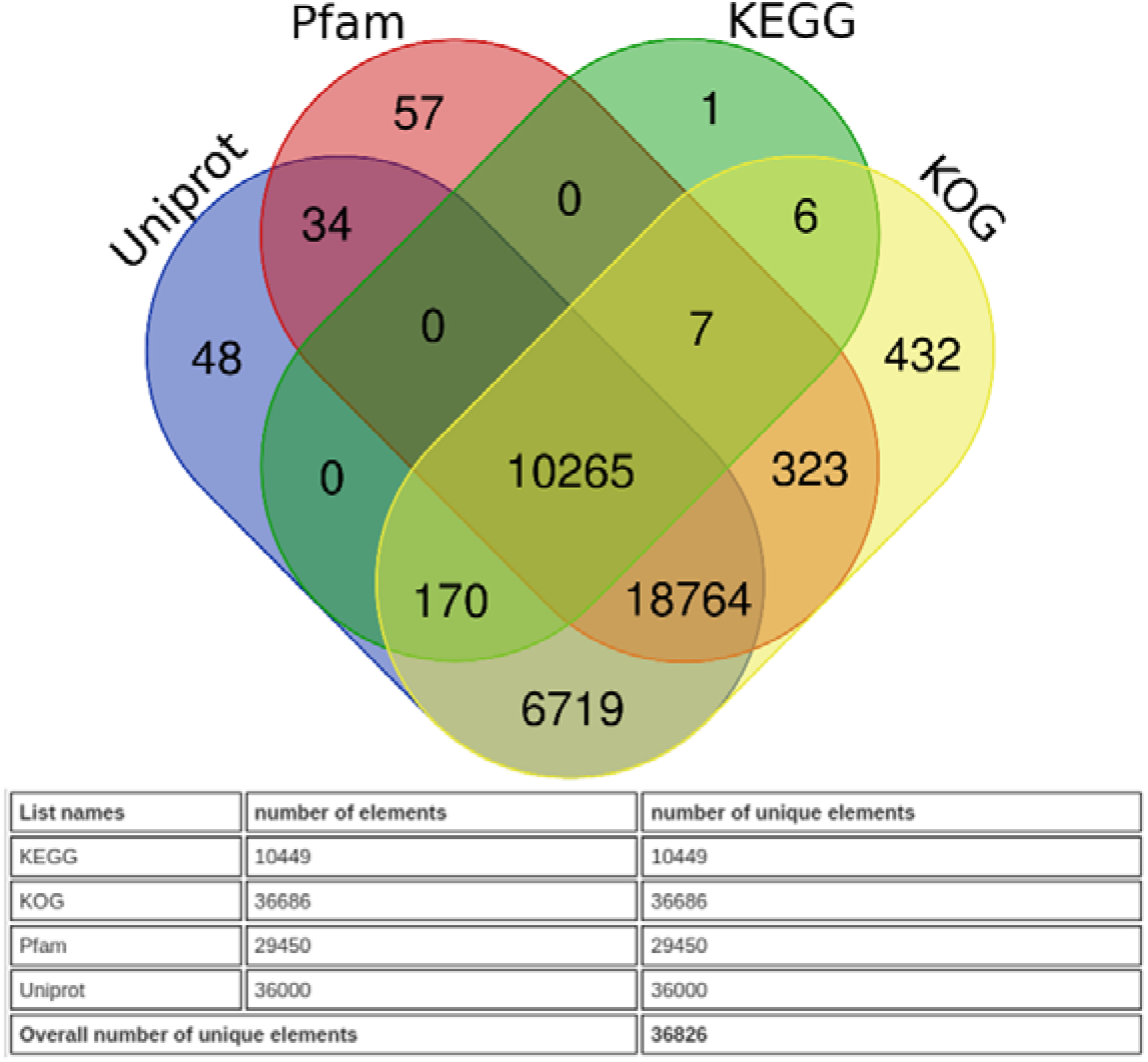
Annotation of predicted coding sequences in the genome of moth bean with various databases

### Gene expression profile

We developed a gene expression profile after mapping the filtered RNA-Seq reads of moth bean onto the coding sequences predicted in the genome, using the scripts provided in the trinity software package (https://github.com/trinityrnaseq/trinityrnaseq/wiki). Leaf tissue sample was used as a reference and the final analysis with analyze_diff_expr.pl script from the trinity software package was done with the following parameters: --matrix RSEM.isoform.TMM.EXPR.matrix -P 0.0001 -C 4 --samples MoB_sampl_desc.txt -- max_DE_genes_per_comparison 40, to get the results for the top 40 hits for the differentially expressed genes in each comparison.

The profile that was developed had a number of features that provide an insight into the developmental processes in the moth bean plant. The expression profile shows clearly demarcated blocks that correspond to genes specifically upregulated in each tissue and represented as blocks (Figure 3a and b). We annotated the differentially expressed genes (DEGs) in each block to obtain information about the major biological processes during moth bean development (Supplementary Table S1). We observed that genes like WAT1 related and Casparian strip integrity factor (CIF1) are integral to root development in moth bean, along with genes encoding germin like protein, cytochrome P450 and MLP like protein.

**Figure 3.**
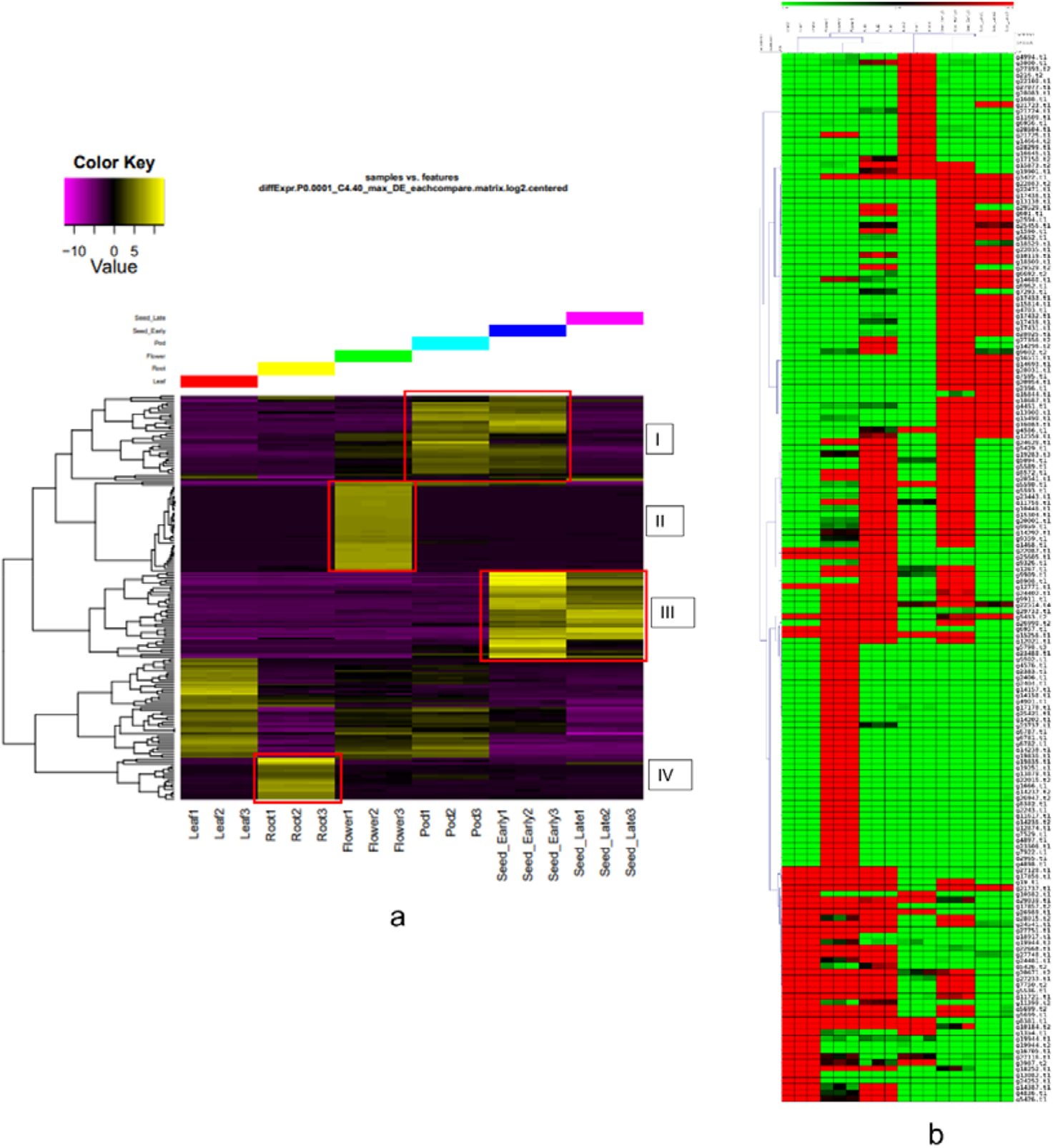
Differential expression profile for various tissues of Moth bean. (a) A representative expression profile for the highly up-regulated genes in tissues of moth bean. The blocks represent genes specifically upregulated in the tissue; block I= Pod and early stages of developing seeds, blockII= flower, blockIII= developing seed, blockIV= root. (b) The expression profile of highly expressed genes after Hierarchical clustering on the MeV software (https://sourceforge.net/projects/mev-tm4/).

Pectinesterases seem to have a vital role in flower development in moth bean, in addition to genes encoding Beta-galactosidase, Chalcone synthase, MYB like transcription factors and L-ascorbate oxidase. We noted the upregulation of genes encoding peroxidases, sugar transporters, proline rich cell wall protein and B3 domain containing protein in both young pod tissue and seeds during early stages of development. The seed tissue, both at early as well as late stages of development, had an abundance of seed storage proteins like glycinin, vicilin, albumin, along with some late embryogenesis abundant (LEA) proteins as well as Abscisic acid insensitive 5 (ABI5) like protein.

### Identification of simple sequence repeats

We used the Microsatellite identification tool (MISA, Beier et al., 2017) to identify the simple sequence repeats (SSRs) in the genome of moth bean with the following parameters: type of repeats/number: mononucleotide/10, dinucleotide/8, trinucleotide/6, tetranucleotide/6, pentanucleotide/3, hexanucleotide/3. The analysis gave us a total of 72043 SSRs with mononucleotide repeats being the most abundant (88.4%) followed by pentanucleotide repeats (7.3%) (Table 3).

**Table 3.** Simple sequence repeats identified in the moth bean genome

|  |  |
| --- | --- |
| Total number of sequences examined | 340 |
| Total size of examined sequences (bp) | 409317563 |
| Total number of identified SSRs | 72043 |
| Number of SSR containing sequences | 28 |
| Number of sequences containing more than 1 SSR | 25 |
| Number of SSRs present in compound formation | 3879 |
| **Distribution to different repeat type classes** |  |
| Unit size | Number of SSRs |
| 1 | 63745 |
| 2 | 662 |
| 3 | 342 |
| 4 | 6 |
| 5 | 5275 |
| 6 | 2013 |

## Discussion

India is the largest producer of moth bean (*Vigna aconitifolia*), a nutritious, underutilised legume crop that is also climate resilient (Ba & Vs, 2020). The grains are rich in protein and carbohydrates and are a popular ingredient in several Indian snacks. Despite the various nutritional and economic prospects, research that focuses on the genomics of this legume crop are limited. This study was conducted with the aim to develop whole genome assembly of moth bean, in an attempt to enrich the genomic resources and to identify genes that are involved in the overall development of the plant.

Moth bean genome is reported to be a diploid genome (2n=22) (Yundaeng et al., 2019) and our k-mer based analysis puts the genome size at about 380Mb. Using the sequence data produced on two high throughput sequencing platforms, we were able to assemble about 409 Mb of genome sequence into 340 scaffolds. Out of these, 13 scaffolds accounted for more than 97% of the total sequence, giving us a high quality first draft of the moth bean genome. This is reinforced by the N50 value of about 30 Mb, which is a good value for a genome assembly and is also supported by the high percentages of single copy orthologs represented in this genome. This implies that the genome assembly has scope for improvement by applying an additional set of softwares in the pipeline to obtain a chromosome-level assembly. The amount of repetitive sequences varies between 40% to 50% in other related *Vigna* species (Pootakham et al., 2021). We noticed a higher percentage of repeat elements in *Vigna aconitifolia*, an observation that can be explored further by comparing the assembly with those of other *Vigna* species, to identify the evolutionary basis for the unique qualities of moth bean.

We identified more than 36,000 protein coding sequences in the genome of moth bean, which is higher as compared to the results of a similar analysis in black gram (*Vigna mungo*) (Pootakham et al., 2021) and mung bean (*Vigna radiata*) (Ha et al., 2021). We developed a tissue specific gene expression profile for moth bean after mapping the RNA-seq reads from leaf, root, flower, young pod and developing seed tissue onto the CDS predicted from the whole genome. The top 40 most highly expressed coding sequences in each tissue give us an insight into the important molecular processes in moth bean development. Most genes identified in this analysis have documented roles in respective tissue development. For example, pectinesterases have been reported to be integral to the process of petal development during flowering (Shirasawa et al., 2022). Similarly, seed storage proteins are expected to be over-represented in seed tissue. However, we also observed genes encoding proline rich cell wall protein to be up regulated in young pods of moth bean, suggesting their role in pod setting and subsequent seed yield. In addition, a gene encoding WAT1 related protein was found to be upregulated in developing seeds of moth bean. This gene has been implicated to have significant influence on seed weight and yield (Pal et al., 2021). In addition, we have identified more than 70,000 SSRs in the genome of moth bean, thereby providing a source of viable molecular markers that can be exploited further for marker-assisted selection. Therefore, this first report of a whole genome assembly for moth bean will form the foundation for future attempts at crop improvement in *Vigna* species.

## Conclusion

The draft of the *Vigna aconitifolia* genome provides considerable genetic insight and data for future crop improvement endeavours. This library of genes that has enormous importance presents a lane to upgrade traits via gene editing and cultivators will be able to cultivate this legume crop variety with type possessing untapped nutritional potential for extensive consumption which has remained underdeveloped throughout centuries. As a result, this study will likely bring nutritionally advantageous legume crops into main stream commercial markets to serve varied targets such as trait improvement in molecular level, thus responding to widespread cases of micronutrient malnutrition around the globe. Therefore, the outcomes of this study address the genetic value of moth bean to be escalated through breeding using allied and applied strategies.

## Ackowledgements

The seeds of moth bean were obtained with the help of the ICAR-CAZRI, Jodhpur, India. The study was part of the national network program on Genetic Enhancement of Minor Pulses (BT/Ag/Network/Pulses-I/2017-2018), supported by the Department of Biotechnology, Govt. of India and we have necessary permission to collect and utilize the seeds for the research purpose.

## Author contributions

SS collected tissue samples from the *V. aconitifolia* var. RMO-435 seeds provided by RKK after thorough screening. SS conducted experimental data generation and visualization. SP designed experiments and analyzed bioinformatics data. ND and SP oversaw the research work. Editing and manuscript writing were conducted by ND, SP, and SS.

## Ethics approval and consent to participate

We confirm that experimental research and field studies are in compliance with relevant institutional, national, and international guidelines and legislation.

## Consent for publication

Not applicable.

