## Supplementary data for "*De novo* assembly of the whole genome of Moth bean (*Vigna aconitifolia*), an underutilized *Vigna* species of India"

| Scaffold ID | Length in bp |
|---|---|
| scaffold_1 | 58823228 |
| scaffold_2 | 47390431 |
| scaffold_3 | 43322040 |
| scaffold_4 | 34354353 |
| scaffold_5 | 30655968 |
| scaffold_6 | 28786593 |
| scaffold_7 | 28462927 |
| scaffold_8 | 26991739 |
| scaffold_9 | 23235433 |
| scaffold_10 | 23092988 |
| scaffold_11 | 22865861 |
| scaffold_12 | 13629529 |
| scaffold_13 | 12032279 |
| scaffold_14 | 1955864 |
| scaffold_15 | 1016720 |
| scaffold_16 | 740118 |
| scaffold_17 | 648579 |
| scaffold_18 | 414031 |
| scaffold_19 | 256279 |
| scaffold_20 | 189634 |
| scaffold_21 | 134954 |
| scaffold_22 | 123011 |
| scaffold_23 | 118275 |
| scaffold_24 | 99832 |
| scaffold_25 | 98798 |
| scaffold_26 | 96931 |
| scaffold_27 | 88062 |
| scaffold_28 | 84477 |
| scaffold_29 | 84094 |
| scaffold_30 | 84090 |
| scaffold_31 | 77887 |
| scaffold_32 | 74853 |
| scaffold_33 | 74621 |
| scaffold_34 | 74426 |
| scaffold_35 | 72661 |
| scaffold_36 | 72439 |
| scaffold_37 | 70557 |
| scaffold_38 | 70238 |
| scaffold_39 | 69162 |
| scaffold_40 | 68956 |
| scaffold_41 | 68725 |
| scaffold_42 | 66835 |
| scaffold_43 | 64654 |
| scaffold_44 | 64319 |
| scaffold_45 | 63548 |
| scaffold_46 | 62318 |
| scaffold_47 | 62258 |
| scaffold_48 | 61617 |
| scaffold_49 | 60854 |
| scaffold_50 | 61231 |
| scaffold_51 | 60689 |

| | |
|---|---|
| scaffold_52 | 61003 |
| scaffold_53 | 60987 |
| scaffold_54 | 59781 |
| scaffold_55 | 59657 |
| scaffold_56 | 59274 |
| scaffold_57 | 58315 |
| scaffold_58 | 58676 |
| scaffold_59 | 58211 |
| scaffold_60 | 55610 |
| scaffold_61 | 54869 |
| scaffold_62 | 54811 |
| scaffold_63 | 54346 |
| scaffold_64 | 53841 |
| scaffold_65 | 50765 |
| scaffold_66 | 49988 |
| scaffold_67 | 49316 |
| scaffold_68 | 48417 |
| scaffold_69 | 48395 |
| scaffold_70 | 48461 |
| scaffold_71 | 48398 |
| scaffold_72 | 47751 |
| scaffold_73 | 47353 |
| scaffold_74 | 47590 |
| scaffold_75 | 46427 |
| scaffold_76 | 45628 |
| scaffold_77 | 44759 |
| scaffold_78 | 44911 |
| scaffold_79 | 44202 |
| scaffold_80 | 43864 |
| scaffold_81 | 43848 |
| scaffold_82 | 43376 |
| scaffold_83 | 43148 |
| scaffold_84 | 43008 |
| scaffold_85 | 42600 |
| scaffold_86 | 42060 |
| scaffold_87 | 41499 |
| scaffold_88 | 41355 |
| scaffold_89 | 41848 |
| scaffold_90 | 41742 |
| scaffold_91 | 41208 |
| scaffold_92 | 41022 |
| scaffold_93 | 40883 |
| scaffold_94 | 40332 |
| scaffold_95 | 40582 |
| scaffold_96 | 40370 |
| scaffold_97 | 39596 |
| scaffold_98 | 39337 |
| scaffold_99 | 39791 |
| scaffold_100 | 39075 |
| scaffold_101 | 39395 |
| scaffold_102 | 38762 |
| scaffold_103 | 39602 |

| | |
|---|---|
| scaffold_104 | 38537 |
| scaffold_105 | 38413 |
| scaffold_106 | 38411 |
| scaffold_107 | 38245 |
| scaffold_108 | 38422 |
| scaffold_109 | 37919 |
| scaffold_110 | 37782 |
| scaffold_111 | 36817 |
| scaffold_112 | 36867 |
| scaffold_113 | 36328 |
| scaffold_114 | 36323 |
| scaffold_115 | 36255 |
| scaffold_116 | 36254 |
| scaffold_117 | 35863 |
| scaffold_118 | 35853 |
| scaffold_119 | 35774 |
| scaffold_120 | 35539 |
| scaffold_121 | 35517 |
| scaffold_122 | 35503 |
| scaffold_123 | 35369 |
| scaffold_124 | 35282 |
| scaffold_125 | 34906 |
| scaffold_126 | 34901 |
| scaffold_127 | 34773 |
| scaffold_128 | 35240 |
| scaffold_129 | 34691 |
| scaffold_130 | 35144 |
| scaffold_131 | 34512 |
| scaffold_132 | 34332 |
| scaffold_133 | 34321 |
| scaffold_134 | 34259 |
| scaffold_135 | 33997 |
| scaffold_136 | 33883 |
| scaffold_137 | 33629 |
| scaffold_138 | 33997 |
| scaffold_139 | 33394 |
| scaffold_140 | 32815 |
| scaffold_141 | 32814 |
| scaffold_142 | 32489 |
| scaffold_143 | 32282 |
| scaffold_144 | 32147 |
| scaffold_145 | 32067 |
| scaffold_146 | 32006 |
| scaffold_147 | 32208 |
| scaffold_148 | 31402 |
| scaffold_149 | 31241 |
| scaffold_150 | 31179 |
| scaffold_151 | 31031 |
| scaffold_152 | 31029 |
| scaffold_153 | 30879 |
| scaffold_154 | 30830 |
| scaffold_155 | 30671 |

| | |
|---|---|
| scaffold_156 | 31148 |
| scaffold_157 | 30174 |
| scaffold_158 | 30141 |
| scaffold_159 | 30110 |
| scaffold_160 | 30089 |
| scaffold_161 | 30039 |
| scaffold_162 | 29938 |
| scaffold_163 | 29892 |
| scaffold_164 | 30346 |
| scaffold_165 | 29793 |
| scaffold_166 | 29406 |
| scaffold_167 | 29371 |
| scaffold_168 | 28953 |
| scaffold_169 | 28951 |
| scaffold_170 | 28905 |
| scaffold_171 | 28672 |
| scaffold_172 | 28565 |
| scaffold_173 | 28482 |
| scaffold_174 | 28267 |
| scaffold_175 | 28168 |
| scaffold_176 | 28164 |
| scaffold_177 | 28029 |
| scaffold_178 | 27663 |
| scaffold_179 | 27647 |
| scaffold_180 | 27644 |
| scaffold_181 | 27368 |
| scaffold_182 | 27355 |
| scaffold_183 | 27294 |
| scaffold_184 | 27275 |
| scaffold_185 | 27165 |
| scaffold_186 | 27162 |
| scaffold_187 | 27025 |
| scaffold_188 | 26689 |
| scaffold_189 | 26474 |
| scaffold_190 | 26428 |
| scaffold_191 | 26130 |
| scaffold_192 | 26080 |
| scaffold_193 | 25796 |
| scaffold_194 | 25648 |
| scaffold_195 | 25515 |
| scaffold_196 | 25483 |
| scaffold_197 | 25371 |
| scaffold_198 | 25288 |
| scaffold_199 | 25215 |
| scaffold_200 | 25206 |
| scaffold_201 | 25632 |
| scaffold_202 | 25117 |
| scaffold_203 | 25094 |
| scaffold_204 | 25058 |
| scaffold_205 | 24938 |
| scaffold_206 | 24834 |
| scaffold_207 | 24633 |

| | |
|---|---|
| scaffold_208 | 25032 |
| scaffold_209 | 24520 |
| scaffold_210 | 24485 |
| scaffold_211 | 24297 |
| scaffold_212 | 24222 |
| scaffold_213 | 24163 |
| scaffold_214 | 24155 |
| scaffold_215 | 24075 |
| scaffold_216 | 24056 |
| scaffold_217 | 24052 |
| scaffold_218 | 23939 |
| scaffold_219 | 23779 |
| scaffold_220 | 23708 |
| scaffold_221 | 23510 |
| scaffold_222 | 23292 |
| scaffold_223 | 23219 |
| scaffold_224 | 23203 |
| scaffold_225 | 23045 |
| scaffold_226 | 22794 |
| scaffold_227 | 22782 |
| scaffold_228 | 22726 |
| scaffold_229 | 22690 |
| scaffold_230 | 22689 |
| scaffold_231 | 22526 |
| scaffold_232 | 22454 |
| scaffold_233 | 22382 |
| scaffold_234 | 22289 |
| scaffold_235 | 22059 |
| scaffold_236 | 22039 |
| scaffold_237 | 22386 |
| scaffold_238 | 21616 |
| scaffold_239 | 21562 |
| scaffold_240 | 21203 |
| scaffold_241 | 20844 |
| scaffold_242 | 20832 |
| scaffold_243 | 20727 |
| scaffold_244 | 20682 |
| scaffold_245 | 20493 |
| scaffold_246 | 20479 |
| scaffold_247 | 20426 |
| scaffold_248 | 20404 |
| scaffold_249 | 20383 |
| scaffold_250 | 20353 |
| scaffold_251 | 20311 |
| scaffold_252 | 20305 |
| scaffold_253 | 20236 |
| scaffold_254 | 20173 |
| scaffold_255 | 20092 |
| scaffold_256 | 20084 |
| scaffold_257 | 19804 |
| scaffold_258 | 19687 |
| scaffold_259 | 19648 |

| | |
|---|---|
| scaffold_260 | 19616 |
| scaffold_261 | 19525 |
| scaffold_262 | 19421 |
| scaffold_263 | 19154 |
| scaffold_264 | 19079 |
| scaffold_265 | 18904 |
| scaffold_266 | 18652 |
| scaffold_267 | 18644 |
| scaffold_268 | 18462 |
| scaffold_269 | 18359 |
| scaffold_270 | 18352 |
| scaffold_271 | 18171 |
| scaffold_272 | 18148 |
| scaffold_273 | 17962 |
| scaffold_274 | 17885 |
| scaffold_275 | 17790 |
| scaffold_276 | 17510 |
| scaffold_277 | 17477 |
| scaffold_278 | 17329 |
| scaffold_279 | 17279 |
| scaffold_280 | 17129 |
| scaffold_281 | 16906 |
| scaffold_282 | 16895 |
| scaffold_283 | 16748 |
| scaffold_284 | 16703 |
| scaffold_285 | 16593 |
| scaffold_286 | 16561 |
| scaffold_287 | 16535 |
| scaffold_288 | 16455 |
| scaffold_289 | 16429 |
| scaffold_290 | 16117 |
| scaffold_291 | 16116 |
| scaffold_292 | 16041 |
| scaffold_293 | 15979 |
| scaffold_294 | 15753 |
| scaffold_295 | 15721 |
| scaffold_296 | 15634 |
| scaffold_297 | 15606 |
| scaffold_298 | 15594 |
| scaffold_299 | 15457 |
| scaffold_300 | 15448 |
| scaffold_301 | 15353 |
| scaffold_302 | 15180 |
| scaffold_303 | 15013 |
| scaffold_304 | 14926 |
| scaffold_305 | 14809 |
| scaffold_306 | 14722 |
| scaffold_307 | 14588 |
| scaffold_308 | 14562 |
| scaffold_309 | 14424 |
| scaffold_310 | 13826 |
| scaffold_311 | 13788 |

| | |
|---|---|
| scaffold_312 | 13611 |
| scaffold_313 | 13543 |
| scaffold_314 | 13314 |
| scaffold_315 | 12841 |
| scaffold_316 | 12760 |
| scaffold_317 | 12545 |
| scaffold_318 | 12497 |
| scaffold_319 | 12289 |
| scaffold_320 | 12224 |
| scaffold_321 | 12186 |
| scaffold_322 | 12104 |
| scaffold_323 | 11846 |
| scaffold_324 | 11695 |
| scaffold_325 | 11676 |
| scaffold_326 | 11622 |
| scaffold_327 | 11236 |
| scaffold_328 | 11193 |
| scaffold_329 | 10951 |
| scaffold_330 | 10424 |
| scaffold_331 | 10313 |
| scaffold_332 | 10307 |
| scaffold_333 | 9876 |
| scaffold_334 | 9835 |
| scaffold_335 | 9386 |
| scaffold_336 | 9326 |
| scaffold_337 | 8994 |
| scaffold_338 | 8661 |
| scaffold_339 | 7976 |
| scaffold_340 | 7907 |

The n50 value is :: 30655968

The N50 Index value is ::5

The total count of contigs is :: 340

The total count of bases is :: 409317563

The avg size of contig is :: 1203875.18529412

largest length of contig is :: 58823228

Smallest length of the contig is :: 7907
